# Self-similar synchronization of calcium and membrane potential transitions during AP cycles predict HR across species

**DOI:** 10.1101/2020.10.26.355412

**Authors:** Syevda Tagirova Sirenko, Kenta Tsutsui, Kirill Tarasov, Dongmei Yang, Ashley N Wirth, Victor A. Maltsev, Bruce D. Ziman, Yael Yaniv, Edward G. Lakatta

## Abstract

**Background:** Translation of knowledge of sinoatrial nodal “SAN” automaticity gleaned from animal studies to human dysrhythmias, e.g. “Sick Sinus” Syndrome (SSS) requiring electronic pacemaker insertion has been sub-optimal, largely because heart rate (HR) varies widely across species.

**Objectives:** To discover regulatory universal mechanisms of normal automaticity in SAN pacemaker cells that are self-similar across species.

**Method:** Sub-cellular Ca^2+^ releases, whole cell AP-induced Ca^2+^ transients and APs were recorded in isolated mouse, guinea-pig, rabbit and human SAN cells. Parametric Ca^2+^ and Vm Kinetic Transitions (PCVKT) during phases of AP cycles from their ignition to recovery were quantified.

**Results:** Although both action potential cycle lengths (APCL) and PCVKT during AP cycles differed across species by ten-fold, trans-species scaling of PCVKT during AP cycles and scaling, of PCVKT to APCL in cells *in vitro*, EKG RR intervals *in vivo*, and BM were self-similar (obeyed power laws) across species. Thus, APCL *in vitro*, HR *in vivo*, and BM of **any** species can be predicted by PCVKT during AP cycles in SAN cells measured in any single species *in vitro*.

**Conclusions:** In designing optimal HR to match widely different BM and energy requirements from mice to humans, nature did not “reinvent pacemaker cell wheels”, but differentially scaled kinetics of gears that regulate the rates at which the “wheels spin”. This discovery will facilitate the development of novel pharmalogic therapies and biologic pacemakers featuring a normal, wide-range rate regulation in animal models and the translation of these to humans to target recalcitrant human SSS.

**Condensed Abstract:** Studies in animal models are an important facet of cardiac arrhythmia research. Because HR differs by over ten-fold between some animals and humans, translation of knowledge about regulatory mechanisms of SAN normal automaticity gleaned from studies in animal models to target human SSS has been sub-optimal. Our findings demonstrating that trans-species self-similarity of sub-cellular and cellular mechanisms that couple Ca^2+^ to Vm during AP cycles can predict heart rate *in vivo* from mice to humans will inform on the design of novel studies in animal models and facilitate translation of this knowledge to target human disease.

## Introduction

Despite decades of electrophysiology research, sinoatrial node (SAN)-initiated arrhythmias are among the most recalcitrant to pharmacologic therapy, and “Sick-Sinus Syndrome” often requires electronic pacemaker insertion. One reason for these therapeutic shortcomings may be that important molecular and cell discoveries of mechanism that regulate SAN pacemaker cell automaticity are mostly made in non-human mammals (e.g. mice, guinea-pig and rabbit), and are often ignored or sluggishly translated to the bedside, because the average human heart rate differs from that of many of these species by up to ten-fold.

Intracellular Ca^2+^ oscillations are a universal property of excitable cells throughout nature (1,2), and occur in SAN pacemaker cells of both human and non-human species (3-7). Different models have been suggested to explain or describe nature’s universal properties. One of these is that long-range correlations throughout nature that obey power law behavior (indicating self-similarity) are manifestations of self-ordered criticality (8). It has indeed been demonstrated that cell-wide Ca^2+^ signals emerge in mouse heart ventricular myocytes when self-organization of spontaneous local **sub-cellular** calcium events achieves criticality (9). Similarly, spontaneous action potentials (APs) in rabbit SAN cell emerge when self-organization of spontaneous local oscillatory Ca^2+^ releases (LCRs) achieves criticality (3,5,10,11).

Recent studies demonstrate that the rate and rhythm of AP firing of SAN cells are controlled by coordinated kinetic transitions in functions of calcium and surface membrane electrogenic molecules that couple to each other in a feed-forward (non-linear) manner to ignite APs. (10,12,13). These kinetic transitions in chemical (Ca^2+^) and electrical (Vm) functional domains that emerge during AP cycles in cells that fire APs reflect activation-inactivation transition kinetics of molecules that determine AP cycle and AP firing rate and rhythm (14). For example, studies in isolated, single rabbit SAN cells indicate that sub-cellular, spontaneous, LCRs activate inward sodium-calcium exchange (NCX) current,(3,15) to regulate the rate of spontaneous surface membrane diastolic depolarization (DD) culminating in a critical cell-wide event, the rapid AP upstroke (5,10,11,16). Further, differences in spontaneous AP firing rates among these cells are highly correlated with differences in periodicity of LCR signals at baseline and during autonomic receptors stimulation (5,17). Numerical sensitivity analyses reveal that only SAN cell models that generate intercellular calcium oscillations in addition to ion channels are able to reproduce the full range of human heart rates (18).

Even though absolute AP cycle lengths from mouse to human differ markedly, because the coupled-clock system that drives SAN automaticity operates on the principle of criticality (19), we reasoned that transitions in both Ca^2+^ and Vm during AP cycle phases are self-similar (obey a power law) both within each species and across species. We also reasoned that Vm and Ca^2+^ transitions across species are self-similar to AP cycle lengths across species and are self-similar to *in vivo* heart rates across species. Further, because the heart is the only organ in which anatomical and physiological properties have been preserved during mammalian species evolution (20), and because it is already known that heart rates derived from EKG RR intervals manifest long-range, power law correlations with body mass (BM) across a wide range of diverse species (21-24), we also hypothesized that the trans-species self-similarity of pacemaker cell coupled-clock mechanisms extends to BM across species. In other terms, we were in quest of identifying a universal concept linking sub-cellular kinetic transitions within a coupled-clock system intrinsic to pacemaker cells to HR across species that matches HR to energy requirements from mice to humans that differ widely in BM.

To this end we recorded Ca^2+^ signals and Vm in single pacemaker cells isolated from mouse, guinea-pig, rabbit and human hearts in order to determine: whether Ca^2+^ and Vm kinetic transitions during AP cycles were self-similar to each other across species; and whether transitions during AP cycles in SAN cells are self-similar to (i) AP cycles lengths *in vitro*, (ii) HR *in vivo* (EKG RR intervals), and (iii) to BM across species.

## Results

### Kinetic transitions in calcium and membrane potential during AP cycles

Subcellular events in SAN pacemaker cells during spontaneous diastolic depolarization (DD) in isolated rabbit SAN cells have been conceptualized as an ignition phase of AP cycles (10). The time at which this Vm acceleration achieves (∼0.15V/s), has been identified as ignition onset in the Vm domain (Fig. 1A) (10). Ignition onset is linked to the occurrence of oscillatory local Ca^2+^ releases (LCRs) (Fig. 1, Video 1), that undergo self-organized phase transitions (Fig. 1A) (11) (during the diastolic depolarization) into roughly periodic, cell-wide ensemble Ca^2+^ signals, culminating in the generation of an AP, a cell-wide event. The AP induces a cell-wide Ca^2+^ transient that faithfully informs on AP cycle length (5,7,10,11,25).

**Figure 1.**
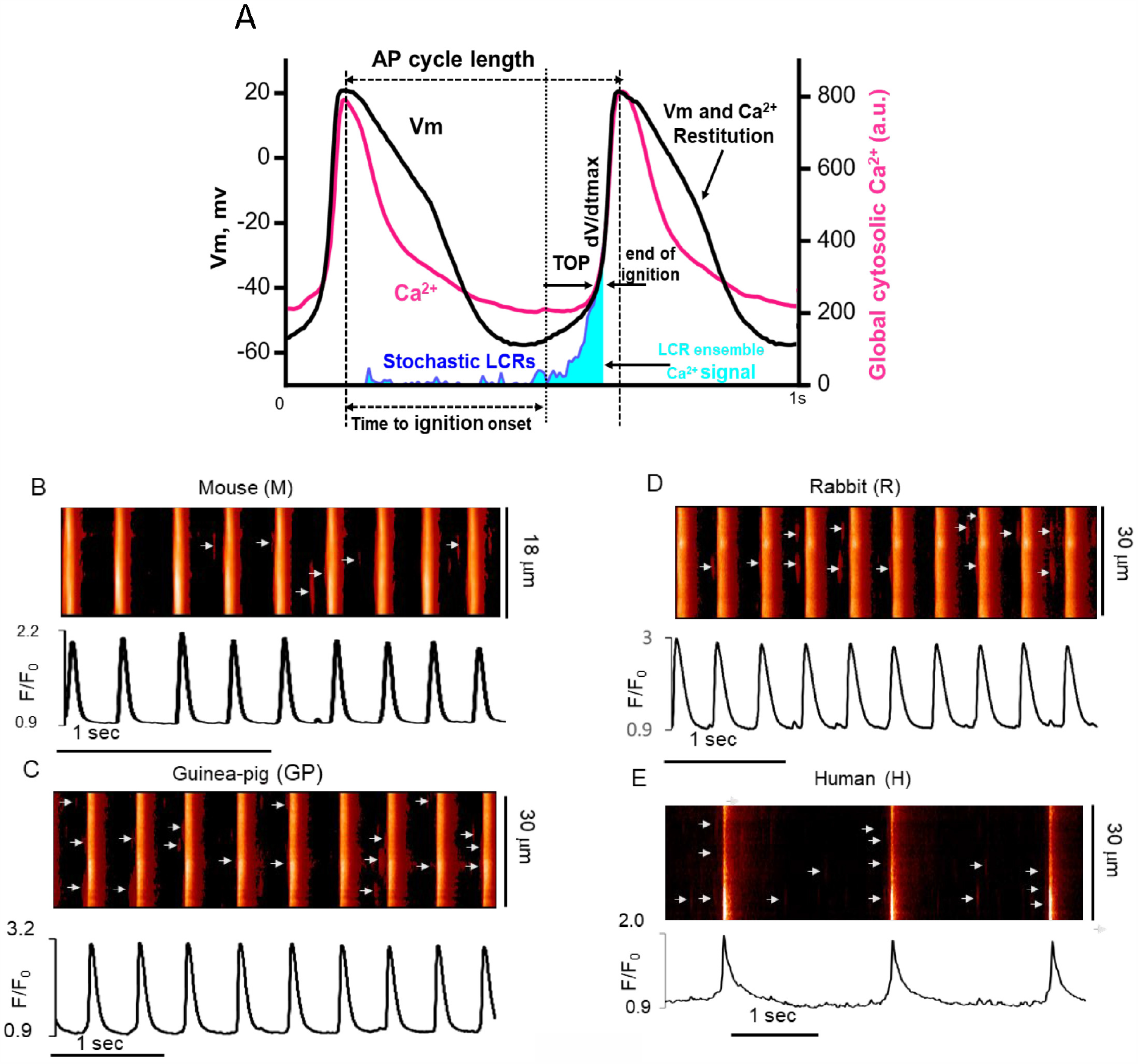
The coupled Ca^2+^ and current oscillator system that drives SAN cells automaticity. (**A**) Simultaneous recordings of Ca^2+^ signals and membrane potential (redrawn from original traces (7)). **(B-D**) Upper panels-representative examples of confocal line-scan images and LCRs (indicated by white arrows on images) in single, spontaneously beating SAN cells; lower panels-AP-induced Ca^2+^ transients (CaT) from the same cells depicted in upper panels, isolated from (**B**) mouse, (**C**) guinea-pig, (**D**) rabbit, (**E**) human hearts and loaded with Ca^2+^ indicator (3-10µM, see methods).

Using 2D imaging (Fig. 1A, Video1 and 2) and confocal microscopy (Fig. 1B-1E), we confirmed that because spontaneous diastolic LCRs, the hallmark diastolic Ca^2+^ signal of the Ca^2+^ clock, are conserved in SAN cells of mice, guinea-pigs, rabbits and human hearts, indicating that a coupled-oscillator system in pacemaker cells is conserved from mice to humans. The completion of the ignition phase in the Vm and Ca^2+^ domain occurs when Vm depolarization and diastolic whole-cell Ca^2+^ transient markedly accelerate, creating the rapid AP and AP-induced Ca^2+^ transient upstroke (Fig. 1A, see methods) (10).

The restitution kinetics of these electrochemical transitions during the AP cycle length are defined in the Vm domain as APD90 (time to 90% AP repolarization) and in the Ca^2+^ domain, as CaT90 (time to 90% decay of the AP-induced calcium transient (Fig. 1A).

### Trans-species self-similarity of kinetic transitions in Ca^2+^ to Vm kinetic transitions during AP cycle

We next established that key kinetic transitions in cells in which Ca^2+^ was measured were self-similar across species, and that Vm kinetic transition in cells in which Vm was measured were self-similar across species (Supplementary Table S2). We next determined whether Ca^2+^ kinetic transitions in SAN cells during AP cycles were self-similar to Vm kinetic transitions (obey power laws) during AP cycles, even though Vm and Ca^2+^ parameters were measured in different SAN cells. Figure 2, Panels A and B demonstrate that the onsets, and completions of the ignition phases and restitution phases (Fig 2, Panel C) in Ca^2+^ and Vm domains are self-similar to each other across species.

**Figure 2.**
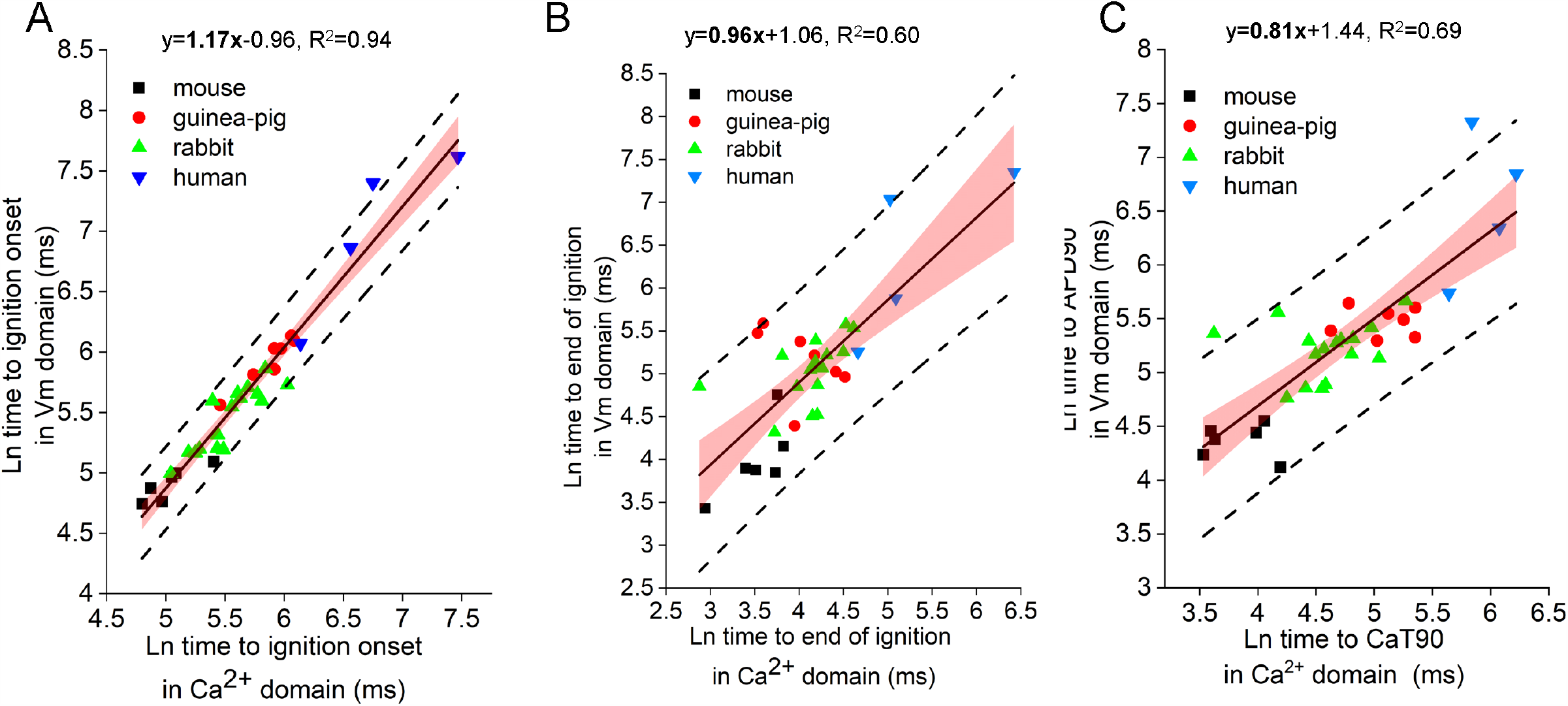
Kinetics of phase transitions in the Vm domain during AP cycles across species are self-similar to kinetics of phase transitions in the Ca^2+^ domain. (**A**) the onset of ignition; (**B**) completion of the ignition phase; (C) restitution. Linear regression of concatenated fit, no weighting; slopes are significantly different from zero (p<0.05). Outside dashed lines-95% prediction band limit; pink - 95% confidence band; n=33 SAN cells for each domain matched by z-scores (see methods); each symbol and color represent a different species.

### Trans-species self-similarity of Ca^2+^ and Vm kinetic transitions during AP cycles to AP cycle lengths

Because clock-coupling determines the AP cycle length, we reasoned that kinetic transitions in both Vm and Ca^2+^ domains during AP cycles across species are self-similar to trans-species AP cycle lengths. Figure 3 shows that this is the case. Thus, although absolute AP cycle lengths differ markedly from mouse to humans, the kinetic transitions in both in Vm and Ca^2+^ domains during ignition and restitution phases of the AP cycle are self-similar to AP cycle lengths across species (Fig. 3 and Supplementary Table S3).

**Figure 3.**
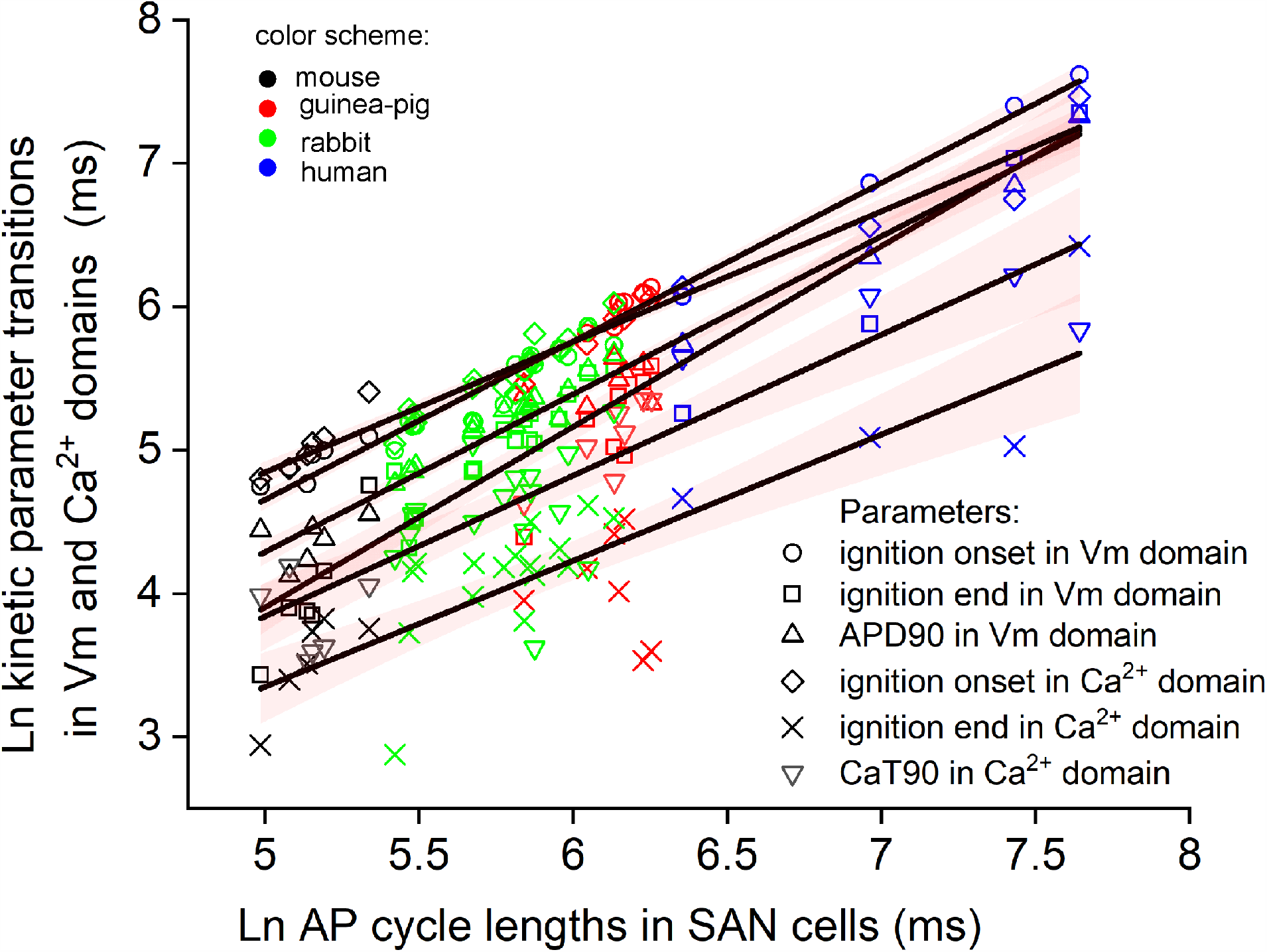
Kinetics transitions during ignition and restitution in Vm and Ca^2+^ domains during AP cycles across species are self-similar to trans-species AP cycle lengths. Linear regression of concatenated fit, no weighting, pink - 95% confidence band; slopes are significantly different from zero (see Supplementary table S3 for regression equations); n=33 SAN cells for Vm and 33 cells for Ca^2+^ domain parameters; each color represents a different species, symbols indicate different parameters. See a Supplementary Table S3 for least square linear regression results.

### Kinetic phase transitions in Vm and Ca^2+^ domains during AP cycles in isolated SAN cells *in vitro* are self-similar to heart rates *in vivo* across species

We next determined whether the self-similarity of ignition and restitution kinetics, and AP firing rate in isolated SAN cells across species *in vitro* (Figs. 1-3) extends to the HRs *in vivo* (EKG RR, cycle lengths). Power law correlations did demonstrate that heart rates (EKG RR intervals) *in vivo* were self-similar to AP cycle ignition in SAN cells *in vitro*, and also to in vitro AP cycle lengths (Figs. 4A and B). Atrial-ventricular conduction times (EKG PR intervals) and ventricular depolarization-repolarization times (EKG QT intervals) were also self-similar across species to AP cycle lengths and AP cycle ignition or restitution intervals in the Vm and Ca^2+^ domains in isolated SAN cells *in vitro* (Fig. 4C and D and Supplementary Table S4). Supplementary Table S4, which list results of linear regression analysis between Vm and Ca^2+^ domains parameters during AP cycles and EKG parameters *in vivo*, indicated that many additional kinetic parameters in single isolated cells and EKG intervals are self-similar across species.

**Figure 4.**
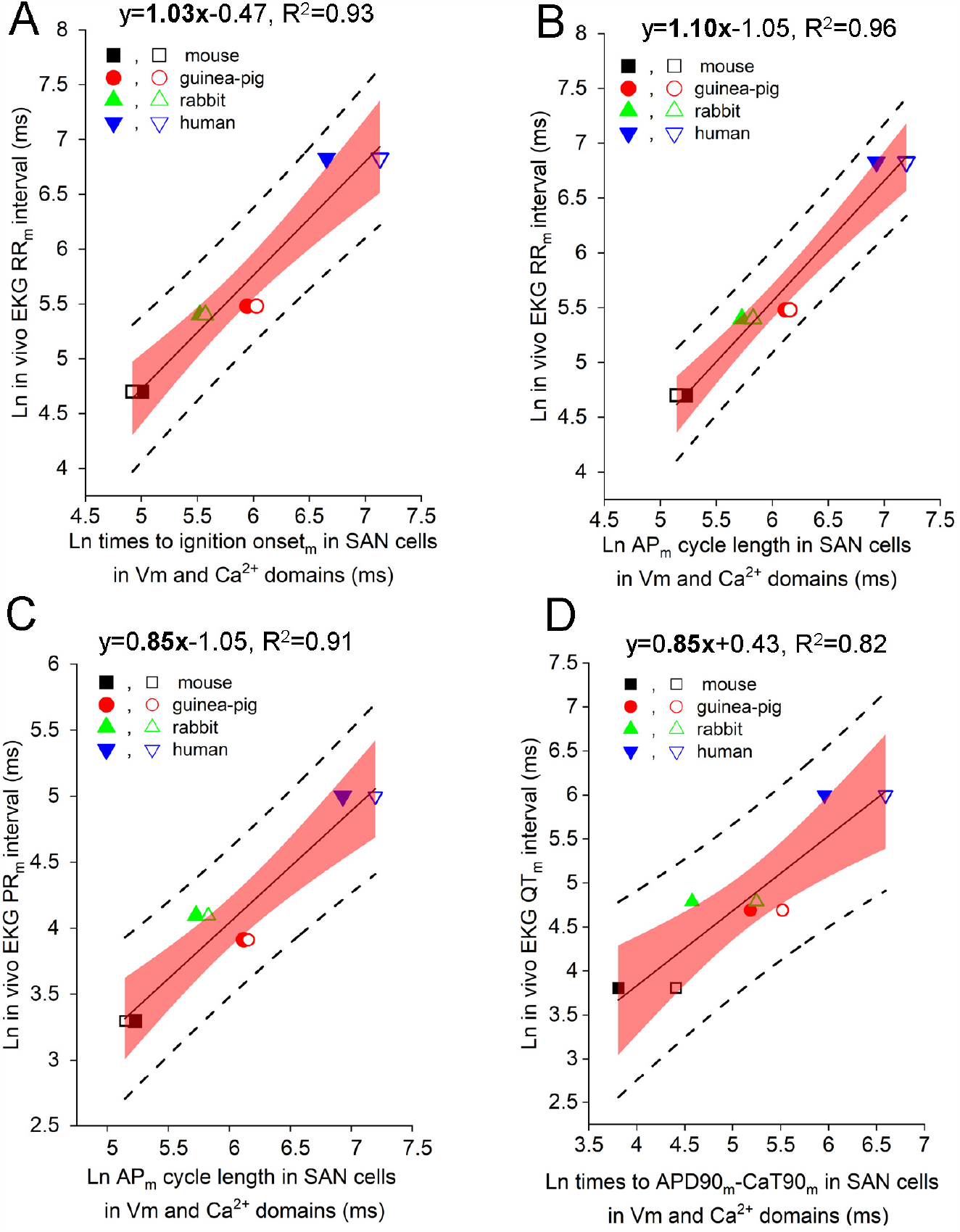
Kinetics of ignition and restitution and AP cycle lengths in Vm and Ca^2+^ domains during AP cycles in isolated SAN cells *in vitro* are self-similar to *in vivo* EKG intervals across species. (**A**) AP cycle lengths in SAN cells *in vitro* vs EKG RR intervals (cycle lengths) *in vivo*; (**B**) AP cycle lengths in SAN cells *in vitro* vs EKG PR intervals (atrioventricular conduction time) *in vivo*; (**C**) times to ignition onset in SAN cells *in vitro* vs EKG PR intervals *in vivo*; (**D**) times to APD90 and CaT90 in SAN cells *in vitro* vs EKG QT intervals (ventricular depolarization-repolarization times) *in vivo*. Open symbols-transmembrane AP (median values) recorded via patch-clamp in Vm domain; closed symbols CaT (median values) recorded via confocal microscopy in Ca^2+^ domain (n=33 SAN cells in each domain). (**A-D**) Linear regression analysis without weighting of values; regression slopes are significantly different from zero (p< 0.05); *in vivo* EKG parameters are taken from published literature (see methods). Outside dashed lines-95% prediction band limit; pink - 95% confidence band. See a Supplementary Table S4 for additional trans-species correlation between EKG intervals in vivo vs SAN cells in vitro.

### Vm and Ca^2+^ domains kinetic parameters during AP cycles in isolated, single SAN cells *in vitro* are self-similar to species body mass (BM)

Because numerous prior studies have demonstrated that HR and BM are self-similar across species (21-24,26-28), and because kinetic parameters in Vm and Ca^2+^ domains measured during AP cycles in single, isolated pacemaker cells *in vitro* and HR *in vivo* manifested self-similarity across species (Fig. 4), we reasoned that kinetic Ca^2+^ and Vm transitions during AP cycles measured in single SAN cells across species *in vitro* must also be self-similar to trans-species BM. Fig 5 revealed that this hypothesis was correct. Specifically, the ignition phase onset (Fig. 5A), end of ignition (Fig. 5B), restitution times (Fig. 5C) of the AP cycle in Vm and Ca^2+^ domains in SAN cells and AP cycle lengths (Fig. 5D) were self-similar to BM across species with nearly the same (0.25) scaling exponent observed previously (22-24,28).

**Figure 5.**
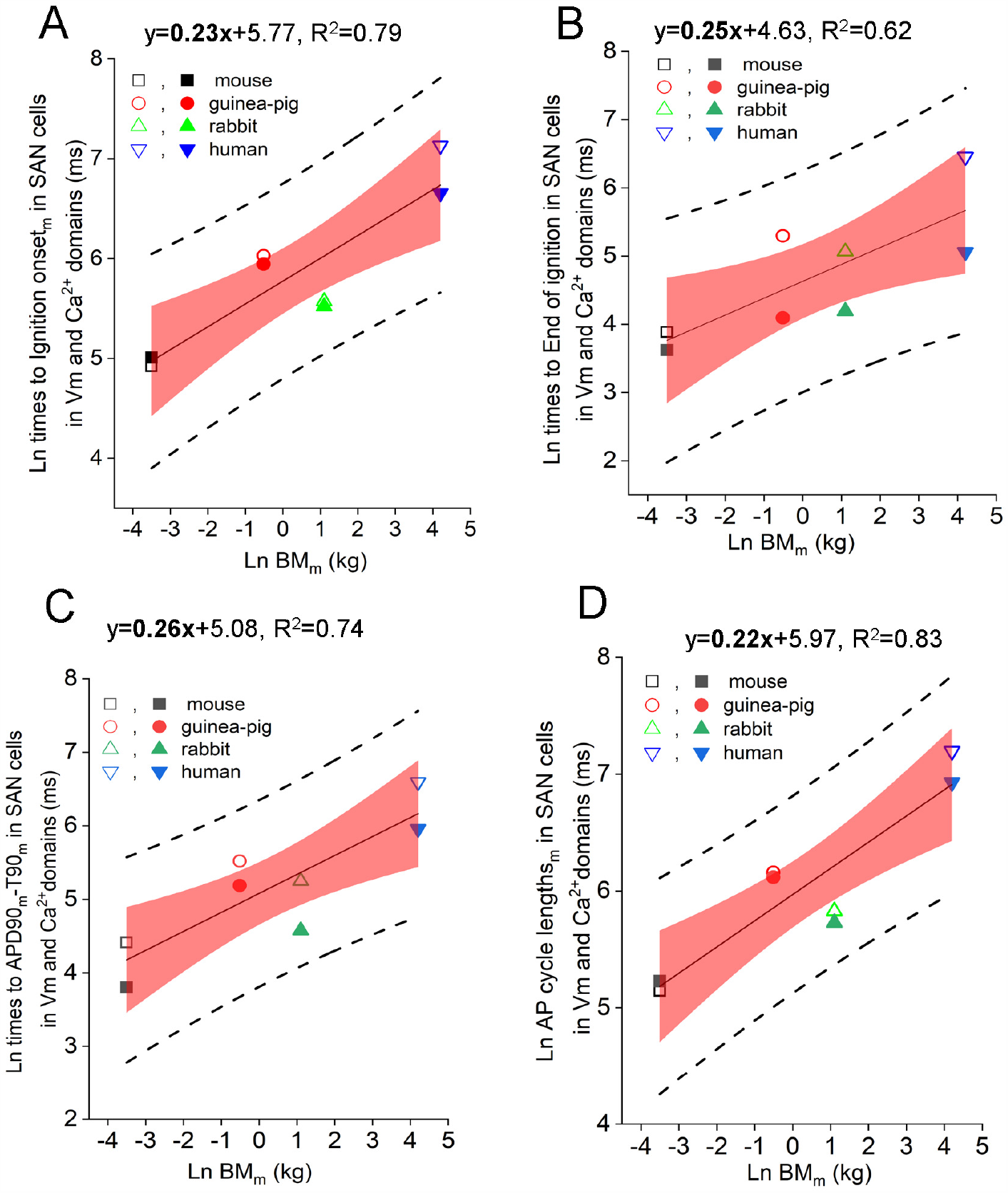
Kinetics of ignition, restitution and AP firing in Vm and Ca^2+^ domains in isolated SAN cells and body mass (BM) are self-similar across species. Median values of BM vs median values of (**A**) Ignition onset (ms), (**B**) End of ignition (ms), (**C**) Cycle lengths (ms) and (**D**) 90% of repolarization (ms) in SAN cells (n=33 for each domain matched by z-scores (see methods)); BM values are taken from published literature (see methods). Open symbols-transmembrane AP recorded via patch-clamp in Vm domain; closed symbols CaT (median values) recorded via confocal microscopy in Ca^2+^ domain. (A-D) Linear regression analysis without weighting of values; slopes are significantly different from zero (p<0.05). Outside dashed lines-95% prediction band limit; pink-95% confidence band; Predicted values from the equations: (**A**) =320.5xBM^0.23±0.04^; (**B**)=102.5xBM^0.25±0.08^; (**C**)=391.5xBM^0.22±0.04^; (**D**)=160.8x BM^0.26±0.06^.

## Discussion

Intracellular Ca^2+^ oscillations are a universal property of excitable cells (1,2) that manifest long-range correlations throughout nature (indicating self-similarity) (21-24,26-28) are manifestations of self-ordered criticality (8,9). Our results uncover novel, long-range (power law) correlations between kinetic transitions of cellular Ca^2+^ and Vm functional domains within AP cycles of single SAN pacemaker cells isolated from hearts across a broad range of mammalian species (Fig. 2, Central Illustration, Video 1). These trans-species power law correlations between cell Ca^2+^ and Vm kinetic transitions during AP cycles extend to: (i) AP cycle lengths of these isolated cells *in vitro*; (ii) *in vivo* heart rates (EKG RR intervals) and other EKG intervals (PR, QT intervals); and (ii) to BM (Figs. 3-5).

Our results also indicate sequence similarity of proteins in Vm and Ca^2+^ domains that regulate pacemaker cell functions (e.g. LCRs, I_NCX_, I_K_ and etc. (Supplementary Table S6)) are highly conserved from mouse to human, and sequences of these proteins are also highly conserved from mouse to human (Supplementary Table S6). Moreover, expression of proteins operative within coupled-oscillator system of pacemaker cells not only pertain to mammals, but are also highly conserved across phyla, from mammals to fishes, and many of these having similar structure (defined by presence of all functional domains in protein sequence) were also identified in insects, flies and worms (Supplementary Table S1). An exception is SCN5A, which is only expressed in mammals and birds (Supplementary Table S1).

Kinetic phase transition in Vm and Ca^2+^ domains, measure during phases of AP cycles in single, isolated SAN cells inform on transitions in activation and inactivation kinetics of molecular functions that underlie the measured functions. Sarcoplasmic reticulum (SR), a Ca^2+^ oscillator, acts as a Ca^2+^ capacitor: pumping Ca^2+^ via a CaATPase (SERCa2), storing Ca^2+^ and releasing Ca^2+^ via ryanodine receptors (RyR) in phosphorylation-dependent manner. The SR Ca^2+^ charge (Ca^2+^ load) is a determinant of the kinetics spontaneous, diastolic local RyR activation that initiates the ignition phase of AP by generating local Ca^2+^ releases (LCRs); Ca^2+^ binding to NCX in the Vm domain generates an inward current (I_NCX_), that accelerates the rate of spontaneous diastolic membrane depolarization. SERCa2, RyR and NCX, activation-inactivation of other molecules (5), in addition to HCN channels (I_f_), Cav 1.3, Cav 3.1 channels (I_CaL-T_) and several K^+^ channels operate in the context a coupled-oscillator system, resulting in feed-forward signaling during the ignition phase, that leads to progressive depolarization of the diastolic membrane potential (3,10,15). In other terms, these proteins are also involved in generation of the electro-chemical gradient oscillations that underlie AP cycles are initiated during the ignition phase of the AP cycle. Criticality is achieved in the Vm domain when the rate of Vm depolarization during an evolving electro-chemical gradient acutely accelerates (the rapid AP uptake) due, in large measure, to activation of Cav 1.2 channels. This is linked to criticality in the Ca^2+^ domain: global activation of RyRs, via a Ca^2+^-induced Ca^2+^ release, resulting in a rapid global increase in cell Ca^2+^, i.e., the AP-induced Ca^2+^ transient. Restitution in the Vm domain occurs as voltage-dependent activation of K^+^ channels effects repolarization of the membrane potential; and in the Ca^2+^ domain, as cytosolic Ca^2+^ decays, due to pumping of a fraction of the Ca^2+^ released into cytosol from SR back to SR (Ca^2+^ recirculation fraction) and extrusion of a fraction of the Ca^2+^ from the cell via NCX (2).

In order to generate the wide range of AP cycle lengths that occur across these species studied, and yet manifest self-similarities of AP cycle lengths and sub-cellular phase transitions across species, coupled-clock molecules (Supplementary Table S6) must be differently “tuned”. This fine-tuning may include differential protein expression levels, differential alternative splicing, and differential post translational modifications e.g. for example differential protein phosphorylation etc. across species. There is dearth of consolidated information regarding fine-tunning of coupled-clock proteins across different species in the literature because of different experimental conditions and experimental design have been employed in different species being compared in different studies.

Available integrated information on species differences in the ion channel current density and channel protein gene expression levels in SAN of different species in the literature is listed in Supplementary Table S7.

In summary, we demonstrate that Ca^2+^ kinetic transitions during AP cycles in SAN cells across species (from mouse to human) are self-similar to trans-species Vm kinetic transitions during AP cycles. Trans-species self-similarity of Ca^2+^ and Vm kinetic transitions during AP cycles in SAN cells *in vitro*, not only predicts the observed self-similarity of AP cycle length in these cells across species, but also predicts trans-species self-similarity of HR *in vivo*. We conceptualized such power law behavior to be a manifestation of trans-species, self-ordered criticality of pacemaker cell molecular functions that regulate HR. Further, because self-similarity of pacemaker cell molecular functions and AP cycle lengths *in vitro* and HR *in vivo* extends to BM explains why EKG RR intervals scale allometrically to BM across species (21-24,26-28). In other terms, in designing appropriate HRs to match BMs and energy requirements of different species from mouse to humans, nature did not “re-invent the wheels” but differentially scaled the gear kinetics that determine the rate at which the “wheels spin”. Together, long-range power law correlations demonstrated here are consistent with the tenet that intra-cellular Ca^2+^ oscillations are a universal property of excitable cells throughout nature (1,2). In order to fulfill the need for a broad range of HRs while protecting against severe bradycardia, nature coupled the criticality mechanism of the Ca^2+^ clock to the limit cycle mechanism of the membrane clock in mice to humans (19). Indeed the full range of human heart rates requires the presence of functional Ca^2+^ clock in addition to membrane clock as demonstrated in single SAN cells (29,30) and in a wide-range sensitivity analyses of numerical models (18).

## Methods

### Ethics statement

The study was performed in accordance with the Guide for the Care and Use of Laboratory Animals published by the National Institutes of Health. The experimental protocols have been approved by the Animal Care and Use Committee of the National Institutes of Health (Protocol # 457-LCS-2021). Adult human hearts not required for transplantation were procured from Washington Regional Transplant Community as previously described (7). None of the donors, age 26-65 years, had a history of major cardiovascular diseases. LVEF: left ventricular ejection fraction. OD, drug overdose, CVA, cerebrovascular accident, CP, cardioplegic. Experimental protocols were approved by the George Washington University Institutional Review Board. Informed donor consents were obtained for all tissue used in this study.

### SAN cells isolation

Single, spontaneously beating SAN cells were isolated from the hearts of adult mice (M), guinea-pigs (GP), rabbits (R) and humans (H) by enzymatic digestions as previously described (6,7,31,32). All methods were performed in accordance with the National Institutes of Health guidelines on human research.

### Transmembrane AP recordings and analyses in SAN cells

Membrane potential was measured in another subset of SAN cells that were not loaded with the Ca^2+^ sensitive indicator. Spontaneous AP were recorded by perforated patch-clamp technique with 0.05 mmol/L of β-escin added to the electrode solution that contained in mmol/L: 120 K-gluconate, 5 NaCl, 5 Mg-ATP, 5 HEPES, 20 KCl, 3 Na_2_ATP (pH adjusted to 7.2 with KOH) (32). SAN cells were continuously superfused with normal Tyrode solution at 35±0.5°C, containing in mmol/L: 140 NaCl, 5.4 KCl, 5 HEPES, 2 MgCl2, 1.8 CaCl2, 5 glucose (pH 7.4). APs were recorded by a standard zero-current-clamp technique (Axopatch 200B, Molecular Devices). APs were corrected for the appropriate liquid junction potential by Clampex 10 Software (Axon Instruments). The AP cycle length and AP characteristics were analyzed via a customized computer program (3,10,33). The AP cycle length was measured as the interval between AP peaks. The program calculated dV/dt (V/s) and the ignition onset (ms) as a time at which Vm dv/dt during diastolic depolarization accelerates to 0.15V/s (10) (see also Fig. 1A). Other measured of AP parameters included: MDP (maximum diastolic potential), TOP (threshold of AP activation when dV/dt reaches 0.5 V/s) (10), end of ignition (time from MDP to take off potential (TOP, ms)) and APD90 (time from AP overshoot to 90% repolarization time, ms).

### Spontaneous diastolic LCRs and AP-induced Ca^2+^ transient (CaT) recordings and analyses in SAN cells

*S*AN cells were loaded with 3-10 µM Ca^2+^ indicator (Cal-520AM or Fluo-4AM) for 10 min, and then were washed with normal Tyrode solution. AP-induced Ca^2+^ transients and spontaneous diastolic local Ca^2+^ releases (LCRs) were recorded in normal Tyrode’s solution (as above) at 35±0.5°C in spontaneously beating SAN cells via confocal microscope. Intracellular Ca^2+^ signals were also recorded using a CMOS 2D camera at 100 FPS (PCO edge 4.2, PCO, Germany) with a conventional inverted microscope and x63 oil objective (Video 2). Our Ca^2+^ transient data include both Fluo-4AM and Cal-520AM measurements. In our earlier studies, we had employed the commonly used Fluo-4AM, but in recent years, and more robust Ca^2+^ probe - Cal-]520AM (AAT Bioquest) with a higher signal/background ratio became available, that we use for detecting Ca^2+^ signals in SAN cells. We did not find a significant difference in mean CaT amplitude between measurements collected with Fluo-4 and Cal-520 AM indicators in SAN cells.

The line-scan mode was executed at a rate 1.92 and 3 ms per scan-line and images were processed with IDL (8.5) software. The scan-line was set along the border of the cell to the specific cell locations beneath the sarcolemma where LCR are present in SAN cells. AP-induced Ca^2+^ transient cycle length (AP cycle length), the faithful proxy of the Vm AP cycle length (25), was defined as the time interval between the peaks of two adjacent AP-triggered Ca^2+^ transients (CaT). Because the use of different indicators could potentially affect the amplitude of CaT, the amplitude of CaTs or spontaneous LCRs was expressed as normalized fluorescence, a peak value (F) normalized to its basal fluorescence (F_0_) rather than just a peak value of (F) in our analysis. Ignition phase onset in Vm was defined previously as the time, when membrane potential accelerates to ∼0.15V/s when the magnitude of inward NCX current begins to increase during diastolic depolarization in SAN cells in which membrane potential and Ca^2+^ were simultaneously measured (10). Ignition phase onset in the Ca^2+^ domain in those studies was a direct read out of the Ca^2+^ signal that prevailed at the Vm ignition onset. In other terms, at ignition onset in the Ca^2+^ domain, the individual LCR Ca^2+^ signals became sufficiently synchronized to generate an ensemble Ca^2+^ signal of sufficient amplitude to accelerate the rate of DD via activation of NCX that generated inward NCX current. In the present study, we used the same acceleration of DD as Lyashkov et al. (10) to define ignition onset in the Vm domain. But since we do not have simultaneous Vm and Ca^2+^ recordings in our study, it is not possible to directly detect the onset of ignition in the Ca^2+^ domain as the calcium signal at the time of ignition onset in the Vm domain. We, therefore, tested various time derivatives of the Ca^2+^ signal during diastole hoping to detect an onset of ignition in the calcium domain. This approach was not satisfactory due to excessive noise in the differentiated Ca^2+^ signal, attributable, in part at least, to LCR occurrence at different times during DD. Therefore, based on our experience in SAN cells and our previous work (10), in order to compare Ca^2+^ domain ignition onset across species, we empirically defined the LCR ensemble ignition onset as the time during DD when the integrated Ca^2+^ signal begins to rise from the background noise and reaches a value 1.5% of the peak value of the subsequent Ca^2+^ transient (see also Supplementary Figure S1). The tight correlation between Ca^2+^ at onset ignition defined as such and ignition onset in Vm (Fig. 2A) indicates the robustness of this approach to define ignition onset in Ca^2+^ domain. Other CaT measurements included: end of ignition (marked as time from baseline to the dCa/dtmax (F/F0/ms^3^)) and restitution of AP-induced CaT, T90 (CaT duration from the peak to 90% of CaT decay, ms) (see Supplementary Figure S1).

### Body mass (BM) and *in vivo* EKG parameters

BM and *in vivo* EKG RR cycle lengths (RR interval), EKG PR intervals (atrioventricular conduction times) and EKG QT intervals (ventricular depolarization and repolarization times) of the diverse species were taken from the literature (23,34-41).

To determine phylogenetic conservation of proteins that regulate pacemaker cell biophysical functions in Mammalian, Birds, Amphibians, Fishes, Insects, Flies and Worms, we used NCBI Ortholog web applet. Corresponding links provided for each compared protein in Supplementary Table S1. Alignments of protein sequences of mouse, guinea-pig rabbit and human proteins were performed with Clustal Omega V.1.2.2 (http://www.clustal.org/omega/) and their accession numbers provided in Supplementary Table S5. Protein sequences were obtained from NCBI database (https://www.ncbi.nlm.nih.gov/gene/) or from UNIPROT (https://www.uniprot.org/). When multiple isoforms were associated with one gene, we selected the longest available or specifically expressed in the heart (if available) for each species for comparison of Mammalian, Birds, Amphibians, Fishes, Insects, Flies and Worms phyla.

### Data processing and statistical analysis

Since Vm and Ca^2+^ were measured in different cells, we computed z-scores of the cycle lengths in Vm and Ca^2+^ domains, separately for each species and then matched the two data sets using R function-matchit (42). Vm (AP) and CaT parameters measured in SAN cells *in vitro* data were transformed to a natural logarithm and plotted as means or as medians of data in individual cells on double logarithmic plots versus means or medians of other Ca^2+^ LCR or Vm parameters or vs medians of species BM and EKG intervals *in vivo*.

Least squares linear regression (no weighting) was applied to determine slopes and 95% confidence and prediction limits using Origin 9.0 software. Slope statistics were tested by ANOVA and t-tests for coefficients. Akaike weights and F-test were used (Origin 9.0) to compare slope differences for multiple datasets; P<0.05 was considered statistically significant. Pearson’s correlation test was used to check the correlation between parameters.

## Abbreviation

AP: action potential
APCL: action potential cycle length
BM: body mass
CaT: AP-induced Ca^2+^ transient
DD: diastolic depolarization
EKG: electrocardiogram
HR: heart rate
LCRs: local sub-cellular Ca^2+^ releases
NCX: sodium-calcium exchanger
RyR: ryanodine receptors
SAN: sinoatrial node
Vm: surface membrane potential

## Author contributions

S.T.S. performed all animal experiments, analyzed the data, wrote and edited the paper; K.T. performed and analyzed human SAN cells experiments; K.V.T. performed phylogenetic comparison across species; D.Y. modified the IDL program to measure the Ca^2+^ ignition onset; A.N.W. provided a 2D video of simultaneous CaT and AP. V.A. provided a program for AP analysis and editing MS; B.D.Z. isolated SAN cells; Y.Y. contributed to discussion and editing of the paper; E.G.L. conceptualized the project, interpreted, wrote and edited the manuscript; all authors commented on the manuscript.

## Acknowledgements

The authors thank Dr. C. Morrell for assistance with statistical analysis and Loretta Lakatta for editorial assistance. This paper was printed in bioRxiv, doi: https://doi.org/10.1101/2020.10.26.355412

## Figure Legends

**Central Illustration. Self-similar synchronization of calcium and membrane potential transitions during AP cycles predict HR across species. (A)** Schematic representation of the experimental setup used to study single SAN cells(top), and original images of LCRs, CaTs or APs recorded by confocal microscopy and patch clamp technique in cells of this study (bottom). **(B)** Ca2+ and Vm kinetic transitions measured during AP cycles in single SAN cells in Ca2+ and VM domains. **(C)** Resting HR varies across species and is negatively correlate to BM. **(D)** Self-similar synchronization of Ca2+ and Vm transitions during AP cycles predict HR across species. Top: long-range power law correlation between the onsets of ignition in the Vm and Ca2+ domains. Bottom: self-similarity between EKG RR intervals in vivo and AP cycle ignition in SAN cells. AP=action potential; BM=body mass; CaT= action potential-induced Ca2+ transient; EKG=electrocardiogram; HR=heart rate; LCRs=spontaneous local sub-cellular calcium releases; RR interval (heart rate in vivo); SAN=sinoatrial node; Vm=membrane potential.

**Online Video 1. Representative example of human SAN cells recorded by a high-speed 2D camera. (A)** 2D image of an isolated, single human SAN cell loaded with Ca^2+^ indicator and recorded by a high speed 2D camera. Bright areas of local fluorescence within the cell represent spontaneous local sub-cellular calcium releases (LCRs). **(B)** Global cytosolic Ca^2+^ (pink) and LCRs ensemble Ca^2+^ signal (cyan blue) from the same human SAN cell.

**Online Video 2. Simultaneous Vm, CaT and LCRs recorded by a high speed 2D camera in rabbit SAN cell. (A)** 2D image of an isolated, single rabbit SAN cell loaded with Ca^2+^ indicator and recorded by a high speed 2D camera. Bright areas of fluorescence within the cell represent sub-cellular LCRs. **(B)** Simultaneous recordings of Ca^2+^ signals, spontaneous local sub-cellular calcium releases (LCRs) and action potentials from the same cell.

---

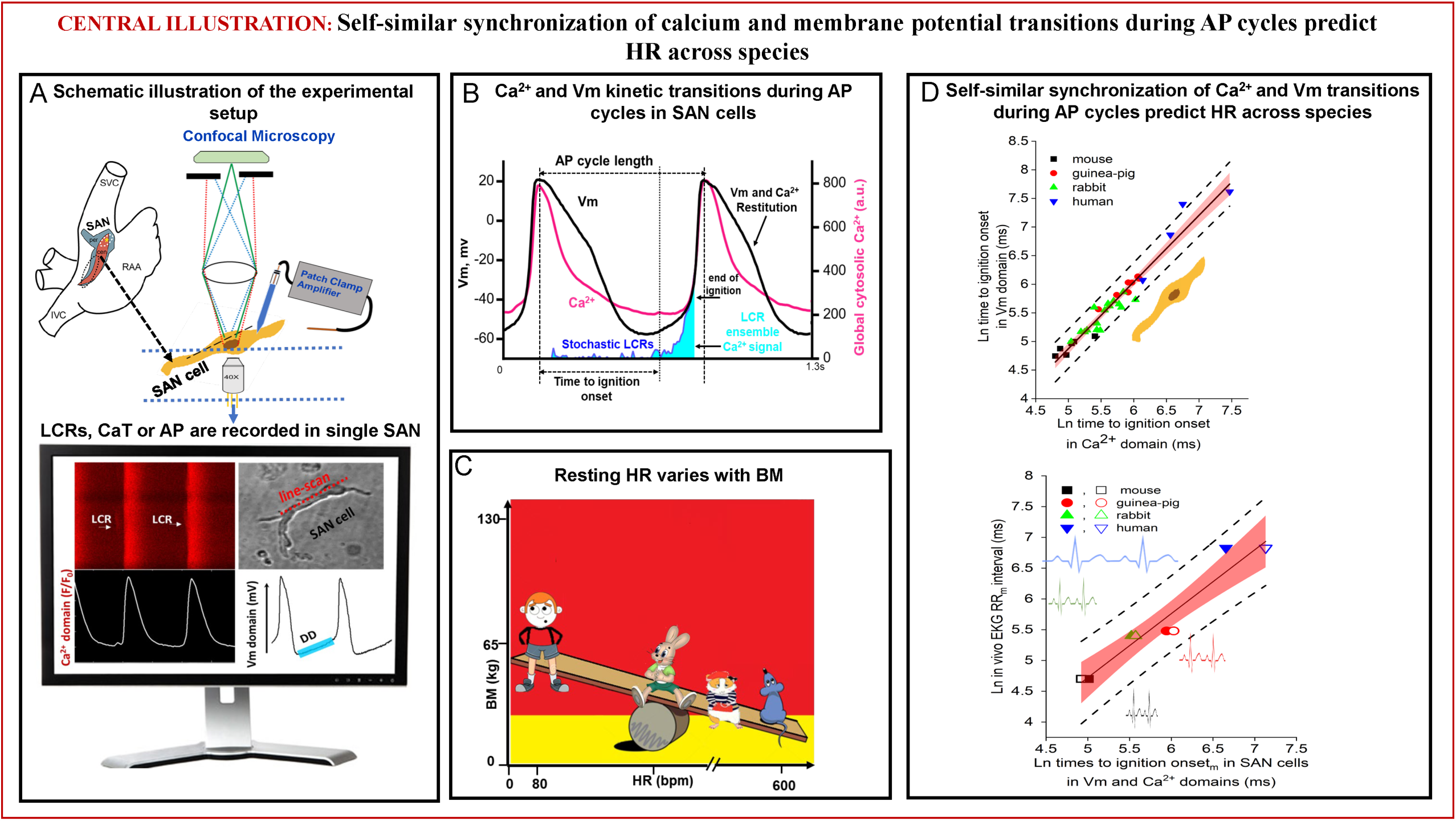

## References

1. Berridge MJ, Lipp P, Bootman MD. The versatility and universality of calcium signalling. Nat Rev Mol Cell Biol 2000;1:11–21.

2. Bers DM. Excitation-contraction coupling and cardiac contractile force. 2nd ed.: Norwell, Mass: Kluwer Academic Publishers; 2001.

3. Bogdanov KY, Maltsev VA, Vinogradova TM, et al. Membrane potential fluctuations resulting from submembrane Ca2+ releases in rabbit sinoatrial nodal cells impart an exponential phase to the late diastolic depolarization that controls their chronotropic state. Circ Res 2006;99:979–87.

4. Chen B, Wu Y, Mohler PJ, Anderson ME, Song LS. Local control of Ca^2+^-induced Ca^2+^ release in mouse sinoatrial node cells. J Mol Cell Cardiol 2009;47:706–15.

5. Lakatta EG, Maltsev VA, Vinogradova TM. A coupled SYSTEM of intracellular Ca^2+^ clocks and surface membrane voltage clocks controls the timekeeping mechanism of the heart’s pacemaker. Circ Res 2010;106:659–73.

6. Sirenko SG, Yang D, Maltseva LA, Kim MS, Lakatta EG, Maltsev VA. Spontaneous, local diastolic subsarcolemmal calcium releases in single, isolated guinea-pig sinoatrial nodal cells. PLoS One 2017;12:e0185222.

7. Tsutsui K, Monfredi OJ, Sirenko-Tagirova SG, et al. A coupled-clock system drives the automaticity of human sinoatrial nodal pacemaker cells. Sci Signal 2018;11.

8. Bak P. How Nature Works: the science of self-organized criticality. New York: Springer-Verlag; 1999.

9. Nivala M, Ko CY, Nivala M, Weiss JN, Qu Z. Criticality in intracellular calcium signaling in cardiac myocytes. Biophys J 2012;102:2433–42.

10. Lyashkov AE, Behar J, Lakatta EG, Yaniv Y, Maltsev VA. Positive Feedback Mechanisms among Local Ca Releases, NCX, and ICaL Ignite Pacemaker Action Potentials. Biophys J 2018;114:1176–89.

11. Maltsev AV, Maltsev VA, Mikheev M, et al. Synchronization of stochastic Ca(2)(+) release units creates a rhythmic Ca(2)(+) clock in cardiac pacemaker cells. Biophys J 2011;100:271–83.

12. Yaniv Y, Lyashkov AE, Sirenko S, et al. Stochasticity intrinsic to coupled-clock mechanisms underlies beat-to-beat variability of spontaneous action potential firing in sinoatrial node pacemaker cells. J Mol Cell Cardiol 2014;77:1–10.

13. Yaniv Y, Sirenko S, Ziman BD, Spurgeon HA, Maltsev VA, Lakatta EG. New evidence for coupled clock regulation of the normal automaticity of sinoatrial nodal pacemaker cells: Bradycardic effects of ivabradine are linked to suppression of intracellular Ca cycling. J Mol Cell Cardiol 2013;62C:80–9.

14. Yang D, Lyashkov AE, Morrell CH, et al. Self-similar action potential cycle-to-cycle variability of Ca2+ and current oscillators in cardiac pacemaker cells bioRxiv 2020 (preprint);doi:10.1101/2020.09.01.277756.

15. Bogdanov KY, Vinogradova TM, Lakatta EG. Sinoatrial nodal cell ryanodine receptor and Na^+^-Ca^2+^ exchanger: molecular partners in pacemaker regulation. Circ Res 2001;88:1254–8.

16. Stern MD, Maltseva LA, Juhaszova M, Sollott SJ, Lakatta EG, Maltsev VA. Hierarchical clustering of ryanodine receptors enables emergence of a calcium clock in sinoatrial node cells. J Gen Physiol 2014;143:577–604.

17. Vinogradova TM, Zhou YY, Maltsev V, Lyashkov A, Stern M, Lakatta EG. Rhythmic ryanodine receptor Ca^2+^ releases during diastolic depolarization of sinoatrial pacemaker cells do not require membrane depolarization. Circ Res 2004;94:802–9.

18. Maltsev VA, Lakatta EG. Numerical models based on a minimal set of sarcolemmal electrogenic proteins and an intracellular Ca clock generate robust, flexible, and energy-efficient cardiac pacemaking. J Mol Cell Cardiol 2013;59:181–95.

19. Weiss J, Qu Z. The Sinus Node: Still Mysterious After All These Years. J Am Coll Cardiol EP 2020;doi:10.1016/j.jacep.2020.09.017.

20. Meijler FL, Meijler TD. Archetype, adaptation and the mammalian heart. Neth Heart J 2011;19:142–8.

21. Gunther B, Morgado E. Allometry of ECG waves in mammals. Biol Res 1997;30:167–70.

22. Noujaim SF, Berenfeld O, Kalifa J, et al. Universal scaling law of electrical turbulence in the mammalian heart. Proc Natl Acad Sci U S A 2007;104:20985–9.

23. Noujaim SF, Lucca E, Munoz V, et al. From mouse to whale: a universal scaling relation for the PR Interval of the electrocardiogram of mammals. Circulation 2004;110:2802–8.

24. West GB, Brown JH, Enquist BJ. The fourth dimension of life: fractal geometry and allometric scaling of organisms. Science 1999;284:1677–9.

25. Yaniv Y, Stern MD, Lakatta EG, Maltsev VA. Mechanisms of beat-to-beat regulation of cardiac pacemaker cell function by Ca(2)(+) cycling dynamics. Biophys J 2013;105:1551–61.

26. Rosati B, Dong M, Cheng L, et al. Evolution of ventricular myocyte electrophysiology. Physiol Genomics 2008;35:262–72.

27. Schmidt-Nielsen K. Scaling, why is animal size so important? Cambridge ; New York: Cambridge University Press; 1984. xi, 241 p. p.

28. Behar JA, Rosenberg AA, Shemla O, et al. A Universal Scaling Relation for Defining Power Spectral Bands in Mammalian Heart Rate Variability Analysis. Front Physiol 2018;9:1001.

29. Vinogradova TM, Bogdanov KY, Lakatta EG. Novel perspectives on the beating rate of the heart. Circ Res 2002;91:e3.

30. Vinogradova TM, Lyashkov AE, Zhu W, et al. High basal protein kinase A-dependent phosphorylation drives rhythmic internal Ca^2+^ store oscillations and spontaneous beating of cardiac pacemaker cells. Circ Res 2006;98:505–14.

31. Liu J, Sirenko S, Juhaszova M, et al. A full range of mouse sinoatrial node AP firing rates requires protein kinase A-dependent calcium signaling. J Mol Cell Cardiol 2011;51:730–9.

32. Vinogradova TM, Zhou YY, Bogdanov KY, et al. Sinoatrial node pacemaker activity requires Ca^2+^/calmodulin-dependent protein kinase II activation. Circ Res 2000;87:760–7.

33. Yaniv Y, Maltsev VA, Ziman BD, Spurgeon HA, Lakatta EG. The “Funny” Current (I_f_) Inhibition by Ivabradine at Membrane Potentials Encompassing Spontaneous Depolarization in Pacemaker Cells. Molecules 2012;17:8241–54.

34. Giannico AT, Garcia DAA, Lima L, et al. Determination of Normal Echocardiographic, Electrocardiographic, and Radiographic Cardiac Parameters in the Conscious New Zealand White Rabbit. J Exot Pet Med 2015;24:223–34.

35. Hess P, Rey M, Wanner D, Steiner B, Clozel M. Measurements of blood pressure and electrocardiogram in conscious freely moving guineapigs: a model for screening QT interval prolongation effects. Lab Anim 2007;41:470–80.

36. Kaese S, Verheule S. Cardiac electrophysiology in mice: a matter of size. Front Physiol 2012;3:345.

37. Lord B, Boswood A, Petrie A. Electrocardiography of the normal domestic pet rabbit. Vet Rec 2010;167:961–5.

38. Merentie M, Lipponen JA, Hedman M, et al. Mouse ECG findings in aging, with conduction system affecting drugs and in cardiac pathologies: Development and validation of ECG analysis algorithm in mice. Physiol Rep 2015;3.

39. Palhares DMF, Marcolino MS, Santos TMM, et al. Normal limits of the electrocardiogram derived from a large database of Brazilian primary care patients. BMC Cardiovasc Disord 2017;17:152.

40. Rijnbeek PR, van Herpen G, Bots ML, et al. Normal values of the electrocardiogram for ages 16-90 years. J Electrocardiol 2014;47:914–21.

41. Xing S, Tsaih SW, Yuan R, et al. Genetic influence on electrocardiogram time intervals and heart rate in aging mice. Am J Physiol Heart Circ Physiol 2009;296:H1907–13.

42. Ho DE, Imai K, King G, Stuart EA. MatchIt: Nonparametric Preprocessing for Parametric Causal Inference. Journal of Statistical Software 2011;42.

