## Supplemental tables and figures for "Self-similar synchronization of calcium and membrane potential transitions during AP cycles predict HR across species"

**Brief title:** Calcium signals predict heart rate across species

**Supplementary information**

***Corresponding authors:**

Dr. Edward G. Lakatta, Laboratory of Cardiovascular Science, NIA/NIH, Biomedical Research Center, 251 Bayview Boulevard, Baltimore, Maryland 21224.

Dr. Syevda Tagirova, Laboratory of Cardiovascular Science, NIA/NIH, Biomedical Research Center, 251 Bayview Boulevard, Baltimore, Maryland 21224.

**Supplementary Figure S1.** Illustration and definition of the phase transition parameters in the $Ca^{2+}$ domain.

**Supplementary Table S1.** Phylogeny of molecules that regulate cell pacemaker functions: mammals to worms.

**Supplementary Table S2.** Least square linear regression of trans-species correlations of kinetic parameters **within** (**A**) Vm and (**B**) $Ca^{2+}$ domains in single, isolated SAN cells.

**Supplementary Table S3.** Least square linear regression of trans-species correlations **between** Vm and $Ca^{2+}$ domain kinetic parameters and AP cycle lengths in single, isolated SAN cells

**Supplementary Table S4.** Least square linear regression of trans-species correlations between Vm and $Ca^{2+}$ domain kinetic parameters (median values) in single SAN cells in vitro and EKG intervals in vivo.

**Supplementary Table S5.** NCBI/Uniprot Accession IDs of sequences used for alignments.

**Supplementary Table S6**. Homology of proteins that regulate SAN pacemaker cell functions in mouse, guinea-pig, rabbit and human.

**Supplementary Table S7.** Comparison of SAN ion current channel densities and channel proteins gene expression levels in different species.

**Supplementary** Alignments.pdf will be provide will be provided when MS accepted for revision.

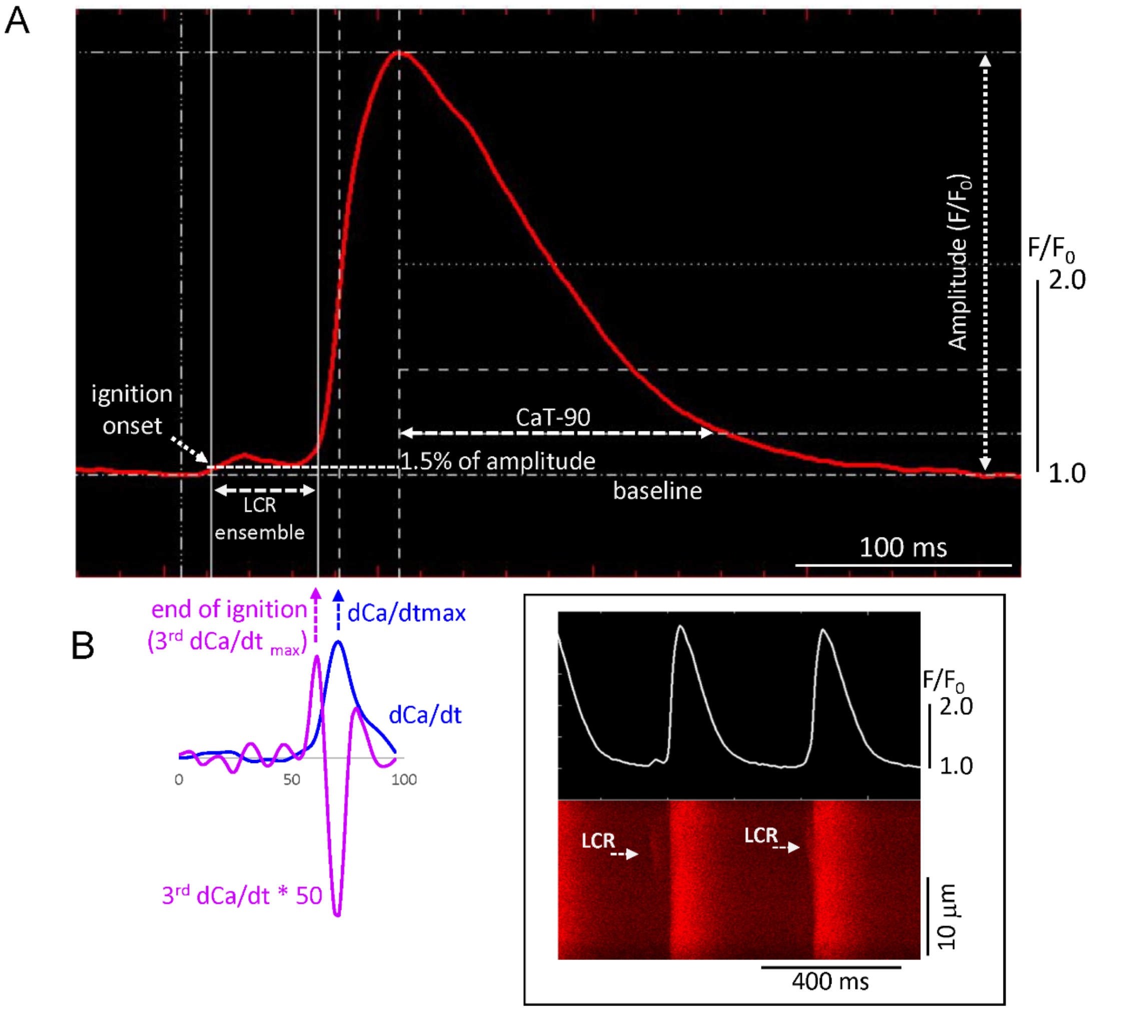


**Supplementary Figure S1.** Illustration and definition of the phase transition parameters in the $Ca^{2+}$ domain. (A) Enlarged AP-induced $Ca^{2+}$ transient (CaT) (2-d CaT trace in B right) defines the phase transition parameters measured with IDL (8.5) software. the ignition onset of LCR ensemble is defined as the time when the integrated $Ca^{2+}$ signal rises from the noise (background) and achieves as a 1.5% of the peak value of the subsequent $Ca^{2+}$ transient amplitude (see also Methods). (**B**) Left: 3d dCa /$dt_{max}$ defines the end of ignition phase, which occurs just prior max rate of rise of CaT. Right: original confocal line-scan image and $Ca^{2+}$ waveform used in **A**.

**S1Table.** Phylogeny of molecules that regulate cell pacemaker functions: mammals to worms.

| Alias | Gene | Mammals | Birds | Amphibians | Fishes | Insects, Flies | Worms | Links to Orthologs | Genes with similar protein architectures |
|---|---|---|---|---|---|---|---|---|---|
| NCX1 | SLC8A1 | + | + | + | + | + | + | https://www.ncbi.nlm.nih.gov/gene/6546/ortholog/?scope=117570 | https://www.ncbi.nlm.nih.gov/gene/6546/ortholog/similargenes/ |
| Hcn2 | HCN2 | + | + | + | + | + | + | https://www.ncbi.nlm.nih.gov/gene/610/ortholog/?scope=117570 | https://www.ncbi.nlm.nih.gov/gene/610/ortholog/similargenes/ |
| Hcn4 | HCN4 | + | + | + | + | + | + | https://www.ncbi.nlm.nih.gov/gene/10021/ortholog/?scope=7776 | https://www.ncbi.nlm.nih.gov/gene/10021/ortholog/similargenes/?scope=33208 |
| Nav1.5 | SCN5A | + | + | | | | | https://www.ncbi.nlm.nih.gov/gene/6331/ortholog/?scope=32524 | https://www.ncbi.nlm.nih.gov/gene/6331/ortholog/similargenes/?scope=7568 |
| Nav1.1 | SCN1A | + | + | + | + | + | | https://www.ncbi.nlm.nih.gov/gene/6323/ortholog/?scope=117570 | https://www.ncbi.nlm.nih.gov/gene/6323/ortholog/similargenes/?scope=6960 |
| Cav 1.2 | CACNA1C | + | + | + | + | | | https://www.ncbi.nlm.nih.gov/gene/775/ortholog/?scope=117570 | https://www.ncbi.nlm.nih.gov/gene/775/ortholog/?scope=117570 |
| Cav1.3 | CACNA1D | + | + | + | + | + | + | https://www.ncbi.nlm.nih.gov/gene/776/ortholog/?scope=7776 | https://www.ncbi.nlm.nih.gov/gene/776/ortholog/similargenes/?scope=33208 |
| Cav3.1 | CACNA1G | + | + | + | + | + | + | https://www.ncbi.nlm.nih.gov/gene/8913/ortholog/?scope=7776 | https://www.ncbi.nlm.nih.gov/gene/8913/ortholog/similargenes/ |
| Kv 7.1 | KCNQ1 | + | + | + | + | + | + | https://www.ncbi.nlm.nih.gov/gene/3784/ortholog/?scope=7776 | https://www.ncbi.nlm.nih.gov/gene/3784/ortholog/similargenes/?scope=33208 |
| MinK | KCNE1 | + | + | + | + | | | https://www.ncbi.nlm.nih.gov/gene/3753/ortholog/?scope=117570 | https://www.ncbi.nlm.nih.gov/gene/3753/ortholog/similargenes/?scope=117570 |
| HERG | KCNH2 | + | + | + | + | + | + | https://www.ncbi.nlm.nih.gov/gene/3757/ortholog/?scope=117570 | https://www.ncbi.nlm.nih.gov/gene/3757/ortholog/similargenes/?scope=2759 |
| KCa1.1 | KCNMA1 | + | + | + | + | + | + | https://www.ncbi.nlm.nih.gov/gene/3778/ortholog/?scope=7776 | https://www.ncbi.nlm.nih.gov/gene/3778/ortholog/similargenes/ |
| KCa2.1 | KCNN1 | + | + | + | + | + | + | https://www.ncbi.nlm.nih.gov/gene/3780/ortholog/?scope=7776 | https://www.ncbi.nlm.nih.gov/gene/3780/ortholog/similargenes/?scope=33213 |
| KCa2.2 | KCNN2 | + | + | + | + | + | + | https://www.ncbi.nlm.nih.gov/gene/3781/ortholog/?scope=7776 | https://www.ncbi.nlm.nih.gov/gene/3781/ortholog/similargenes/?scope=6157 |
| KCa2.3 | KCNN3 | + | + | + | + | | | https://www.ncbi.nlm.nih.gov/gene/3782/ortholog/?scope=7776 | https://www.ncbi.nlm.nih.gov/gene/3782/ortholog/similargenes/?scope=33208 |
| Orai1 | ORAI1 | + | + | + | + | + | + | https://www.ncbi.nlm.nih.gov/gene/84876/ortholog/?scope=7776 | https://www.ncbi.nlm.nih.gov/gene/84876/ortholog/similargenes/?scope=6072 |
| Orai2 | ORAI2 | + | + | + | + | + | + | https://www.ncbi.nlm.nih.gov/gene/80228/ortholog/?scope=117570 | https://www.ncbi.nlm.nih.gov/gene/80228/ortholog/similargenes/?scope=6072 |
| Orai3 | ORAI3 | + | + | + | + | + | + | https://www.ncbi.nlm.nih.gov/gene/93129/ortholog/?scope=117570 | https://www.ncbi.nlm.nih.gov/gene/93129/ortholog/similargenes/?scope=6072 |
| Serca2a | Atp2a2 | + | + | + | + | + | + | https://www.ncbi.nlm.nih.gov/gene/488/ortholog/?scope=7776 | https://www.ncbi.nlm.nih.gov/gene/488/ortholog/similargenes/?scope=33208 |
| RyR2 | RyR2 | + | + | + | + | + | | https://www.ncbi.nlm.nih.gov/gene/6262/ortholog/?scope=7776 | https://www.ncbi.nlm.nih.gov/gene/6262/ortholog/similargenes/?scope=33213 |
| Pln | PLN | + | + | + | + | | | https://www.ncbi.nlm.nih.gov/gene/5350/ortholog/?scope=7898 | https://www.ncbi.nlm.nih.gov/gene/5350/ortholog/similargenes/?scope=7776 |
| Casq2 | CASQ2 | + | + | + | + | | | https://www.ncbi.nlm.nih.gov/gene/845/ortholog/?scope=117570 | https://www.ncbi.nlm.nih.gov/gene/845/ortholog/similargenes/?scope=117570 |
| IP3-kinase / | ITPKA | + | + | + | + | + | + | https://www.ncbi.nlm.nih.gov/gene/3706/ortholog/?scope=7776 | https://www.ncbi.nlm.nih.gov/gene/3706/ortholog/similargenes/?scope=2759 |
| FKBP12 | FKBP1A | + | + | + | + | + | + | https://www.ncbi.nlm.nih.gov/gene/2280/ortholog/?scope=117570 | https://www.ncbi.nlm.nih.gov/gene/2280/ortholog/similargenes/?scope=2759 |
| Stim1 | STIM1 | + | + | + | + | + | + | https://www.ncbi.nlm.nih.gov/gene/6786/ortholog/similargenes/?scope | https://www.ncbi.nlm.nih.gov/gene/6786/ortholog/?scope=7776 |
| Stim2 | STIM2 | + | + | + | + | + | + | https://www.ncbi.nlm.nih.gov/gene/57620/ortholog/?scope=7776 | https://www.ncbi.nlm.nih.gov/gene/57620/ortholog/similargenes/?scope=6072 |
| ADCY1 | ADCY1 | + | + | + | + | + | + | https://www.ncbi.nlm.nih.gov/gene/107/ortholog/?scope=7776 | https://www.ncbi.nlm.nih.gov/gene/107/ortholog/similargenes/?scope=33208 |
| ADCY8 | ADCY8 | + | + | + | + | + | + | https://www.ncbi.nlm.nih.gov/gene/114/ortholog/?scope=7776 | https://www.ncbi.nlm.nih.gov/gene/114/ortholog/similargenes/?scope=33208 |
| PDE4A | PDE4A | + | + | + | + | + | + | https://www.ncbi.nlm.nih.gov/gene/5141/ortholog/?scope=7776 | https://www.ncbi.nlm.nih.gov/gene/5141/ortholog/similargenes/?scope=2759 |

**Supplementary Table S2.** Least square linear regression of trans-species correlations of kinetic parameters **within** (**A**) Vm and (**B**) $Ca^{2+}$ domains in single, isolated SAN cells.

**(A)**

| Vm domain | Ignition onset | | | End of ignition | | | APD90 | | |
|---|---|---|---|---|---|---|---|---|---|
| parameters (ms) | Equation | $R^2$ | p-value | Equation | $R^2$ | p-value | Equation | $R^2$ | p-value |
| Ignition onset | | | | | | | | | |
| end of ignition | y=1.10x-1.18 | 0.85 | <0.05 | | | | | | |
| APD90 | y=0.96x-0.16 | 0.92 | <0.05 | y=0.78x+1.34 | 0.87 | <0.05 | | | |
| Cycle length | y=0.89x+0.90 | 0.98 | <0.05 | y=0.71x+2.32 | 0.89 | <0.05 | y=0.88x+1.28 | 0.96 | <0.05 |

**(B)**

| $Ca^{2+}$ domain | Ignition onset | | | End of ignition | | | APD90 | | |
|---|---|---|---|---|---|---|---|---|---|
| parameters (ms) | Equation | $R^2$ | p-value | Equation | $R^2$ | p-value | Equation | $R^2$ | p-value |
| Ignition onset | | | | | | | | | |
| end of ignition | y=0.96x-1.27 | 0.69 | <0.05 | | | | | | |
| CaT90 | y=1.00x-0.31 | 0.66 | <0.05 | y=0.66x+1.99 | 0.39 | <0.05 | | | |
| Cycle length | y=0.99x+0.90 | 0.99 | <0.05 | y=0.75x+2.79 | 0.76 | <0.05 | y=0.67x+2.74 | 0.68 | <0.05 |

*N=33 SAN cells for Vm and 33 SAN cells for $Ca^{2+}$ domain.*

**Supplementary Table S3.** Least square linear regression of trans-species correlations **between** Vm and $Ca^{2+}$ domain kinetic parameters and AP cycle lengths in single, isolated SAN cells.

| Kinetic transitions variables | Equation | $R^2$ | p-value |
|---|---|---|---|
| **Ignition onset in Vm domain** | y=1.105 x-0.871 | 0.979 | <2E-16 |
| **Ignition end in Vm domain** | y= 1.260 x-2.396 | 0.894 | <2E-16 |
| **APD90 in Vm domain** | y=1.101 x-1.217 | 0.964 | <2E-16 |
| **Ignition onset in $Ca^{2+}$ domain** | y= 0.910 x+0.294 | 0.957 | <2E-16 |
| **Ignition end in $Ca^{2+}$ domain** | y= 0.880 x-1.051 | 0.671 | 5.5E-09 |
| **CaT90 in $Ca^{2+}$ domain** | y=0.982 x-1.072 | 0.738 | 1.60E-10 |

*N=33 SAN cells for Vm and 33 SAN cells for $Ca^{2+}$ domain*

**Supplementary Table S4.** Least square linear regression of trans-species correlations between Vm and $Ca^{2+}$ domain kinetic parameters (median values) in single SAN cells in vitro and EKG intervals in vivo.

| | EKG intervals (ms) in vivo | | | | | | | | |
|---|---|---|---|---|---|---|---|---|---|
| | RR | | | PR | | | QT | | |
| Vm and $Ca^{2+}$ domains (ms) in vitro | Equation | $R^2$ | p-value | Equation | $R^2$ | p-value | Equation | $R^2$ | p-value |
| **Cycle length** | y=1.10x-1.05 | 0.96 | <0.05 | y=0.85x-1.05 | 0.91 | <0.05 | y=1.10x-1.84 | 0.93 | <0.05 |
| **times to Ignition onset** | y=1.03x-0.47 | 0.93 | <0.05 | y=0.80x-0.58 | 0.87 | <0.05 | y=1.03x-1.25 | 0.90 | <0.05 |
| **times to APD90-T90** | y=0.85x+1.23 | 0.84 | <0.05 | y=0.66x+0.69 | 0.80 | <0.05 | y=0.85x+0.43 | 0.82 | <0.05 |
| **times to end of ignition** | y=0.70x+2.24 | 0.65 | <0.05 | y=0.57x+1.48 | 0.65 | <0.05 | y=0.73x+1.38 | 0.67 | <0.05 |

*N=33 SAN cells for Vm and 33 cells for $Ca^{2+}$ domain;* in vivo EKG parameters are taken from published literature (see methods).

**Supplementary Table S5.** NCBI/Uniprot Accession IDs of sequences used for alignments.

| Symbol | mouse | guinea-pig | rabbit | human |
|---|---|---|---|---|
| **SLC8A1** | NP_035536 | NP_001166490 | NP_001164429 | sp\|P32418\|NAC1_HUMAN |
| **HCN2** | O88703 | H0WDP7 | ^ | Q9UL51 |
| **HCN4** | NP_001074661 | XP_003462239 | NP_001076176 | sp\|Q9Y3Q4\|HCN4_HUMAN |
| **SCN5A** | Q9JJV9 | H0V4Z8 | G1STZ7 | Q14524 |
| **SCN1A** | A2APX8 | H0UZ29 | G1SSP8 | P35498 |
| **CACNA1C** | NP_033911 | NP_001166394 | NP_001129994 | sp\|Q13936\|CAC1C_HUMAN |
| **CACNA1D** | NP_001289566 | XP_005008320 | XP_017199370 | NP_000711.1 |
| **CACNA1G** | NP_033913 | XP_013004978 | XP_008269518 | sp\|O43497\|CAC1G_HUMAN |
| **KCNQ1** | P97414 | O70344 | Q9MYS6 | P51787 |
| **KCNE1** | P23299 | Q60409 | Q28705 | P15382 |
| **KCNH2** | O35219 | Q8WNY2 | H0VZT8 | Q12809 |
| **KCNMA1** | Q08460 | A0A286XP95 | Q9BG98 | Q12791 |
| **KCNN1** | Q9EQR3 | H0W7E2 | ^ | Q92952 |
| **KCNN2** | P58390 | H0UUD3 | G1T862 | Q9H2S1 |
| **KCNN3** | P58391 | H0VU03 | A0A5F9C4T7 | Q9UGI6 |
| **ORAI1** | Q8BWG9 | H0W6E9 | G1TJL6 | Q96D31 |
| **ORAI2** | Q8BH10 | A0A286Y0K4 | G1U3B1 | Q96SN7 |
| **ORAI3** | Q6P8G8 | A0A286XUS9 | G1TRG3 | Q9BRQ5 |
| **ATP2A2** | NP_733765 | XP_003462979 | NP_001082790 | NP_001103610 |
| **RyR2** | NP_076357 | XP_023422283 | NP_001076226 | sp\|Q92736\|RYR2_HUMAN |
| **PLN** | NP_001135399 | XP_012998839 | NP_001076090 | sp\|P26678\|PPLA_HUMAN |
| **CASQ2** | O09161 | A0A286Y396 | P31235 | O14958 |
| **ITPKA** | Q8R071 | H0UZF9 | G1TMC6 | P23677 |
| **FKBP1A** | P26883 | A0A286XID4 | P62943 | P62942 |
| **STIM1** | P70302 | A0A286Y154 | G1T594 | Q13586 |
| **STIM2** | P83093 | H0V0T4 | G1T6V7 | Q9P246 |
| **ADCY1** | NP_033752 | XP_005001973 | XP_008260055 | NP_066939 |
| **ADCY8** | NP_033753 | XP_003467396 | XP_002710595 | NP_001106 |
| **PDE4A** | NP_899668 | XP_003460919 | ^ | P27815 |

^ *Not mapped to rabbit genome.*

**Supplementary Table S6.** Homology of proteins that regulate SAN pacemaker cell functions in mouse, guinea-pig, rabbit and human.

| ***Function domains*** | **Alias** | **Gene** | ***Pairwise Identity***[#] |
|---|---|---|---|
| ***Vm domain*** | | | |
| $I_{NCX}$ | Ncx1 | SLC8A1 | 93.8% |
| $I_f$ HCN2 | Hcn2 | HCN2 | 77.5%^ |
| $I_f$ HCN4 | Hcn4 | HCN4 | 88.9% |
| $I_{Nav1.5}$ | Nav1.5 | SCN5A | 88.2% |
| $I_{Nav1.1}$ | Nav1.1 | SCN1A | 97.0% |
| $I_{Ca\text{-}L}$ α1C | Cav 1.2 | CACNA1C | 92.4% |
| $I_{Ca\text{-}L}$ α1D | Cav1.3 | CACNA1D | 96.6% |
| $I_{Ca\text{-}T}$ α1G | Cav3.1 | CACNA1G | 91.6% |
| $I_{Ks}$ α | Kv 7.1 | KCNQ1 | 87.4% |
| $I_{Ks}$ β | MinK | KCNE1 | 74.8% |
| $I_{kr}$ | HERG | KCNH2 | 94.7% |
| $I_{BK1}$ | KCa1.1 | KCNMA1 | 91.3% |
| $I_{SK1}$ | KCa2.1 | KCNN1 | 83.1%^ |
| $I_{SK2}$ | KCa2.2 | KCNN2 | 95.1% |
| $I_{SK3}$ | KCa2.3 | KCNN3 | 94.1% |
| $I_{CRAC}$ | Orai1 | ORAI1 | 89.0% |
| $I_{CRAC}$ | Orai2 | ORAI2 | 94.6% |
| $I_{CRAC}$ | Orai3 | ORAI3 | 89.5% |
| ***$Ca^{2+}$ domain*** | | | |
| CaATPase | Serca2a | ATP2A2 | 98.4% |

| | | | |
|---|---|---|---|
| RyR | RyR2 | RyR2 | 97.0% |
| Phospholamban | Pln | PLN | 98.1% |
| Calsequestrin | Casq2 | CASQ2 | 89.1% |
| Inositol-trisphosphate 3-kinase A | IP3-kinase A | ITPKA | 68.3% |
| FK506 binding protein 12 | FKBP12 | FKBP1A | 98.1% |
| Stromal Interaction Molecule 1 | Stim1 | STIM1 | 91.8% |
| Stromal Interaction Molecule 2 | Stim2 | STIM2 | 91.8% |
| ***cAMP regulators*** | | | |
| Adenylate cyclase 1 | ADCY1 | ADCY1 | 93.7% |
| Adenylate cyclase 8 | ADCY8 | ADCY8 | 97.6% |
| Phosphodiesterase 4A | PDE4A | PDE4A | 81.3%^ |

#*Alignment and sequence acquisition numbers are illustrated in Supplementary Table S6 and Alignments.pdf; ^ not mapped to rabbit genome*

**Supplementary Table S7.** Comparison of ion current channel density and channel proteins gene expression levels in SAN of different species.

| Current/Gene | Current Density (ratio) | | | | Expression level (ratio) | | | |
|---|---|---|---|---|---|---|---|---|
| ***Ignition*** | **M** | **R** | **RB** | **H** | **M** | **R** | **RB** | **H** |
| **$I_f$/HCN4** | 5x vs H | 1.8x vs H | 3.3x vs H | | 7x vs H | 2x vs H | 8x vs H | |
| **$I_{NCX}$/NCX1** | – 4x vs RB | | 1x vs H | | 1.4x vs H | 1x vs H | 5x vs H | |
| **$I_{CaT}$/Cav3.1** | 2x vs RB | | | | 16x vs H | 17x vs H | 1x vs H | |
| **$I_{CaL}$/Cav1.3** | | | | | 1x vs H | 1x vs H | 4x vs H | |
| **INa/Nav1.1-1.4** | | | | | 7.5 x H | 2x vs H | | |
| **RyR** | | | | | 6x vs H | 5x vs H | 1x vs H | |
| **SERCA** | | | | | 70 vs H | 14x vs H | 5x vs H | |
| ***End of Ignition*** | | | | | | | | |
| **$I_{CaL}$/Cav1.2** | 1x vs H | 1x vs H | 1x vs H | | 1x vs H | 1x vs H | 1x vs H | |
| ***Repolarizatiuon*** | | | | | | | | |
| **$I_{kr}$ (activation)/hERG** | 1x vs RB | 1x vs M | | | 1x vs RB | 1x vs M | | |
| **$I_{ks}$/KCNQ1** | | | | | | | | |
| **$I_{BK1}$/KCNMA1** | | | | | | | | |
| **$I_{SK1-3}$/KCNN1-3** | 1x vs RB | | | | | | | |
| **SERCA[&]** | | | | | | | | |

*Data were collected from the literature from Li et al[1]. and other studies[2-7]. M-mouse, R-rat, RB-rabbit, H-human. &see above (ignition). Values that were 1 or <1, we assigned as 1- no difference. Empty cells - no quantitative comparison was found.*

References


1. Li, J., Dobrzynski, H., Lei, M. & Boyett, M. R. Comparison of Ion Channel Gene Expression in the Sinus Node of the Human, Rabbit, Rat and Mouse. Comput Cardiol Conf 43, 1105-1108 (2016).
2. Chen, W. T. et al. Apamin modulates electrophysiological characteristics of the pulmonary vein and the Sinoatrial Node. Eur J Clin Invest 43, 957-963, doi:10.1111/eci.12125 (2013).
3. Clark, R. B. et al. A rapidly activating delayed rectifier K+ current regulates pacemaker activity in adult mouse sinoatrial node cells. Am J Physiol Heart Circ Physiol 286, H1757-1766, doi:10.1152/ajpheart.00753.2003 (2004).
4. Lai, M. H. et al. BK channels regulate sinoatrial node firing rate and cardiac pacing in vivo. Am J Physiol Heart Circ Physiol 307, H1327-1338, doi:10.1152/ajpheart.00354.2014 (2014).
5. Torrente, A. G. et al. Contribution of small conductance K(+) channels to sinoatrial node pacemaker activity: insights from atrial-specific Na(+) /Ca(2+) exchange knockout mice. J Physiol 595, 3847-3865, doi:10.1113/JP274249 (2017).
6. Zimmer, T., Haufe, V. & Blechschmidt, S. Voltage-gated sodium channels in the mammalian heart. Glob Cardiol Sci Pract 2014, 449-463, doi:10.5339/gcsp.2014.58 (2014).
7. Verkerk, A. O., van Borren, M. M. & Wilders, R. Calcium transient and sodium-calcium exchange current in human versus rabbit sinoatrial node pacemaker cells. ScientificWorldJournal 2013, 507872, doi:10.1155/2013/507872 (2013).
